# Differential DNA modification of an enhancer at the *IGF2* locus affects dopamine synthesis in patients with major psychosis

**DOI:** 10.1101/296756

**Authors:** Shraddha Pai, Peipei Li, Bryan Killinger, Lee Marshall, Peixin Jia, Ji Liao, Arturas Petronis, Piroska Szabo, Viviane Labrie

## Abstract

Dopamine dysregulation is central to the pathogenesis of diseases with major psychosis, but its molecular origins are unclear. In an epigenome-wide investigation in neurons, individuals with schizophrenia and bipolar disorder showed reduced DNA modifications at an enhancer in *IGF2*, which disrupted the regulation of the dopamine synthesis enzyme tyrosine hydroxylase and striatal dopamine levels in transgenic mice. Epigenetic control of this enhancer may be an important molecular determinant of psychosis.

## Main Text

Epigenetic misregulation of the genome can trigger long-lasting changes to neurodevelopmental programs, synaptic architecture, and cellular signaling that can increase the risk of psychotic disorders, such as schizophrenia and bipolar disorder^1-3^. In particular, abnormalities in the epigenetic mark DNA modification have been detected in the brain of schizophrenia and bipolar disorder patients, and may play a role disease pathophysiology^4-6^. However, DNA methylation studies of bulk brain tissue are confounded by sample-level variation in the proportion of different cell types. In addition, epigenetic changes occurring within neurons can be masked by the predominant glial signal; there are ~3.6 times more glia than neurons in the human frontal cortex gray matter^7^. In major psychosis, neurons exhibit transcriptional, structural (decreases in dendritic spine density) and neurotransmitter signaling abnormalities. However, whether or not neurons of patients with psychotic illnesses exhibit DNA methylation alterations that contribute to their malfunction has yet to be examined in detail.

We fine-mapped DNA modifications in neuronal nuclei (NeuN+) isolated by flow cytometry from *post-mortem* frontal cortex of the brain of individuals diagnosed with schizophrenia, bipolar disorder, and controls (n=29, 26, and 28 individuals, respectively; Supplementary Table 1). We performed an epigenome-wide association analysis (EWAS) using Illumina MethylationEPIC arrays surveying 812,663 CpG sites (Fig. 1 and Supplementary Fig. 1). After controlling for age, sex, and genetic ancestry, we identified 18 regions with significant DNA modification changes in patients with major psychosis (comb-p Šidák *p* < 0.05; Supplementary Table 2). Differentially methylated regions were enriched in pathways related to embryonic development, synaptic function, and immune cell activation (Fig. 1b, Supplementary Table 3); these pathways were also enriched in transcriptomes from the same tissue samples (Fig. 1b; RNA-sequencing; n=17 cases, 17 controls; Supplementary Table 4-6, and Supplementary Fig. 2a). Pre- and post-natal transcriptional dynamics of genes differentially expressed in psychosis is significantly correlated with those of synaptic development genes (BrainSpan; *p* < 0.001; resampling; Supplementary Fig. 2b). Furthermore, 13 of the 18 differentially-methylated regions demonstrated significant genetic-epigenetic interactions in *cis* (FDR < 0.05; genotypes from Infinium PsychArray-24; 36 of 56 CpG probes in regions; 2,212 of 13,552 tested SNPs) (Fig. 1c, and Supplementary Table 7). Additionally, one differentially-methylated region at the *HLA* locus demonstrated significant genetic-epigenetic interactions with known genetic risk factors for schizophrenia (FDR < 0.05; 4,373 SNPs tested; Supplementary Table 8). Therefore, neurons in major psychosis show significant changes in DNA modifications, some of which are associated with genetic state. These converge on transcriptional changes that affect early development, impair synaptic activity, and raise immune responses.

**Figure 1.**
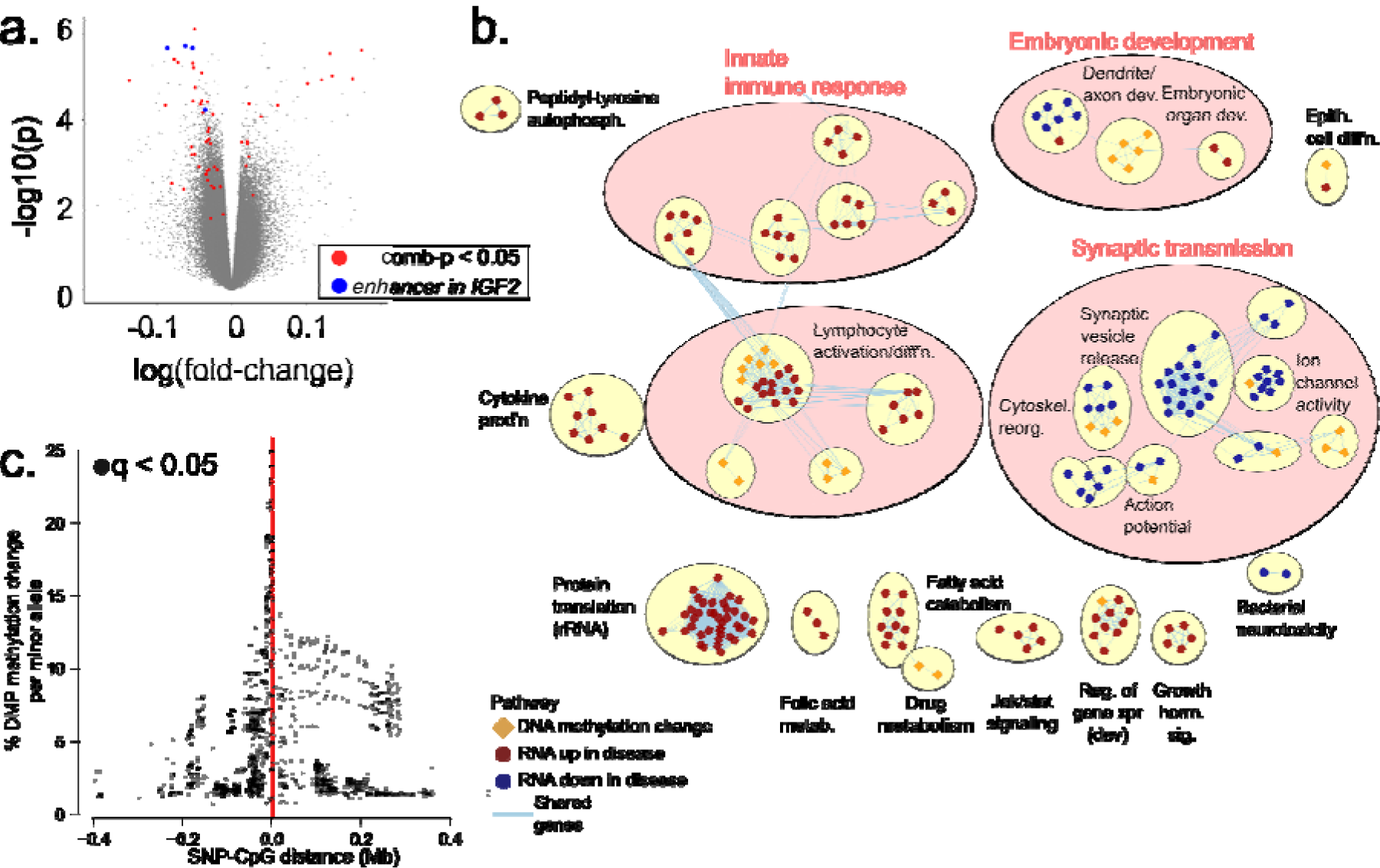
DNA methylation and transcriptional changes in frontal cortex neurons of major psychosis patients converge on synaptic transmission, embryonic development, and immune pathways. (**a**) Volcano plot showing DNA methylation differences in neurons of major psychosis patients (n=45) relative to controls (n=28). Disease-specific DNA methylation changes were identified, after controlling for sex, age, post-mortem interval, and first two genetic principal components. (**b**) Pathways significantly altered in major psychosis by DNA methylation and/or transcription. Nodes show pathways enriched with differentially methylated regions in psychosis (diamonds; q < 0.05, hypergeometric test) or enriched in differentially expressed genes (circles; q < 0.05, GSEA preranked), and edges indicate common genes. Node fills indicate up-(blue) or down-(red) regulation in disease; epigenetic pathways indicate change in disease (orange). Node clusters of similar pathways are grouped in pink circles (Enrichment Map, AutoAnnotate). Clusters with 1-2 nodes are not shown, unless these have both epigenetic and transcriptomic pathways. (**c**) Significant genetic-epigenetic *cis* interactions of differentially-methylated regions. The y-axis shows percent change in DNA methylation with increasing number of minor alleles, (q < 0.05; n=4,762 interactions of 2,212 SNPs with CpGs in 18 DMR regions).

Notably, two of the top differentially modified regions in major psychosis neurons were located at the 3’ end of the *IGF2* gene (Šidák *p* < 10^−3^) (Fig. 2a; Supplementary Table 2a). Both schizophrenia and bipolar patients were consistently hypomethylated at the *IGF2* locus, relative to controls (3-9% mean probe-level hypomethylation; Fig. 2b). Hypomethylation at *IGF2* was even observed in neurons of unmedicated major psychosis patients (Supplementary Fig. 3). The effect of disease on average *IGF2* modification remained significant after controlling for age, sex, PMI, smoking status, medication, and first two genetic principal components (nested model ANOVA, *p* < 0.02; Supplementary Fig. 3), as well as in an analysis limited to individuals of European ancestry (Šidák *p* < 0.001; Supplementary Table 2b). We then fine-mapped DNA modifications at the *IGF2* genomic area (~161 kb) in neurons, using a targeted bisulfite sequencing assay (n=4 cases, 3 controls). This analysis confirmed the significant hypomethylation of the *IGF2* region in neurons of major psychosis cases (*p* < 10^−4^, mixed effects model; Fig. 2c, Supplementary Table 9). We did not find evidence of *cis*-acting genetic-epigenetic effects for any of the probes in the differentially methylated *IGF2* region (FDR > 0.05; Supplementary Table 10). We also did not find evidence that cases and controls differed in allele-specific expression of *IGF2* (Supplementary Fig. 4).

**Figure 2.**
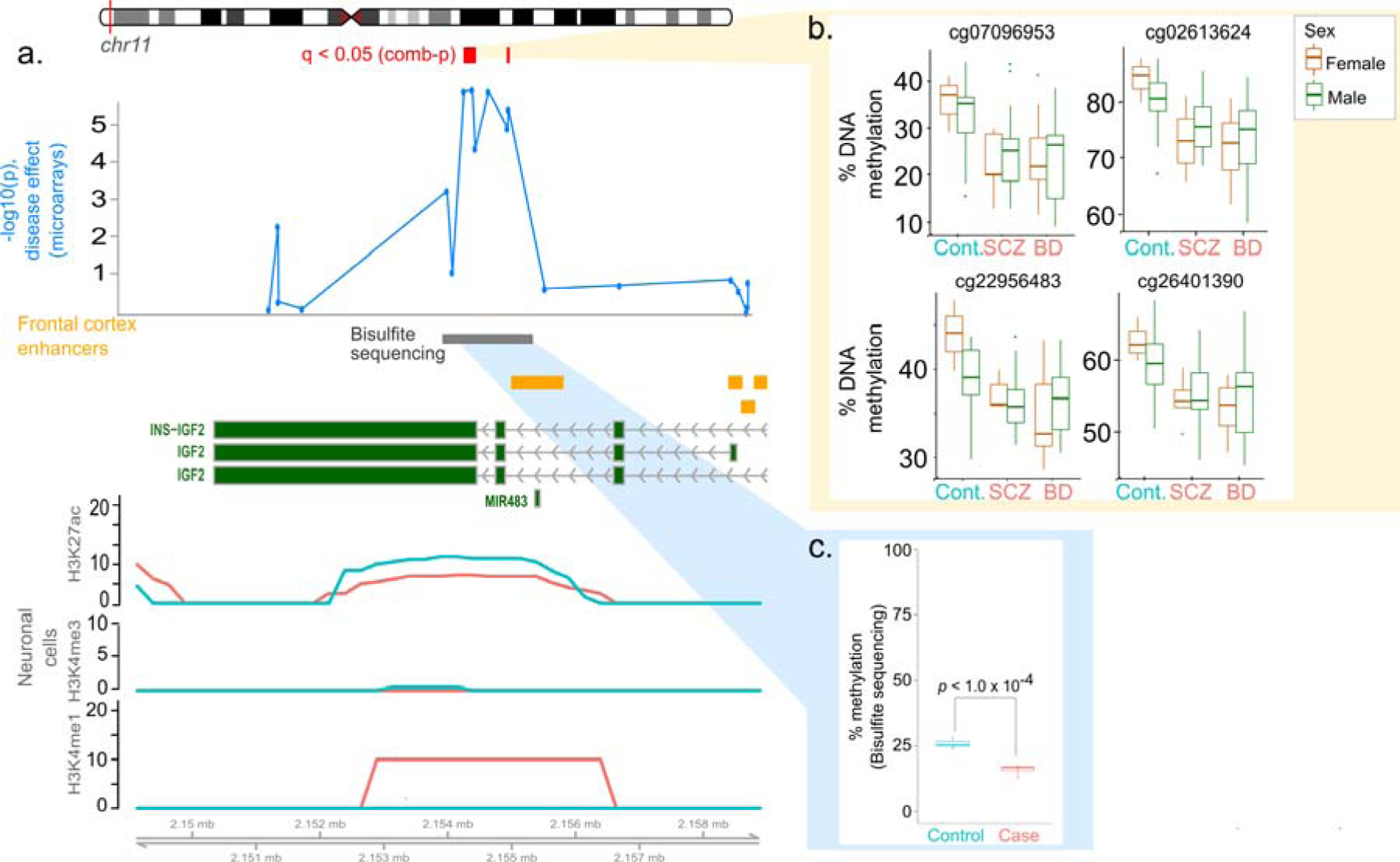
An enhancer at *IGF2* is differentially methylated in neuronal cells in major psychosis. (**a**) Hypomethylation in psychosis at the *IGF2* locus. The view shows differentially methylated regions (red, comb-p) and nominal probe-level p-values (blue) in major psychosis cases (n=45) relative to controls (n=28), as identified by EPIC arrays. Also shown are regulatory elements in the region: adult frontal cortex enhancers (orange; NIH Epigenomics Roadmap), and histone marks in neuronal cells derived from olfactory neuroepithelium obtained from controls (teal) and schizophrenia (salmon) (PsychENCODE; case,control sample sizes: H3K27ac: n=16, n=12; H3K4me3: n=8, n=8; H3K4me1: n=2, n=1). (**b**) CpG probe-level methylation within differentially methylated *IGF2* region, by diagnostic subgroup and sex. (**c**) Validation of *IGF2* hypomethylation using targeted bisulfite sequencing. Average % DNA methylation in a region in *IGF2* that is differentially methylated in major psychosis. Box plots show the % DNA methylation averaged over the ~1.3 kb enhancer region. Bases with ≥10× coverage are included in the analysis. P-value from nested mixed-effects model for effect of disease, after controlling for age and sex (n=4 cases, 3 controls; technical replicates included as random effect). Base-level methylation estimates from EPIC arrays and targeted bisulfite sequencing were strongly correlated (Pearson’s coefficient R=0.57, *p* < 10^−4^).

The hypomethylated *IGF2* locus in major psychosis overlapped an enhancer in the adult frontal cortex (Fig. 2a; data from NIH Roadmap). Furthermore, in neurons derived from olfactory neuroepithelium, histone marks characteristic of enhancers (H3K4me1; H3K27ac) were present in schizophrenia patients, but depleted in controls (data from PsychENCODE; Fig. 2a). Assessment of chromatin interactions in the prefrontal cortex by analysis of Hi-C data revealed that this enhancer targets the tyrosine hydroxylase gene promoter (Fig. 3a). Tyrosine hydroxylase is the rate-limiting enzyme for the production of the neurotransmitter dopamine. Dopamine dysregulation in the cortex and striatum of both patients with schizophrenia and bipolar disorder is centrally involved in the cognitive and psychotic symptoms^8, 9^. Reduced DNA modifications at the *IGF2* enhancer was associated with elevated levels of TH protein levels in the human frontal cortex (*p*<0.05; Fig. 3b), supporting the hypothesis that this enhancer modulates dopamine synthesis. We then examined transgenic mice carrying an intergenic *Igf2* enhancer deletion. In the striatum, inactivation of the *Igf2* enhancer led to a decrease in TH protein levels and in dopamine (*p*<0.05; Fig. 3c and 3d); this effect was not observed in the frontal cortex (Supplementary Fig. 5). These data collectively suggest that in schizophrenia and bipolar disorder, epigenetic disruption of enhancer activity at the *IGF2* locus in neurons leads to abnormalities in dopaminergic signaling. Hypomethylation of the enhancer at *IGF2* may be an important contributor to pathogenesis of psychotic symptoms.

**Figure 3.**
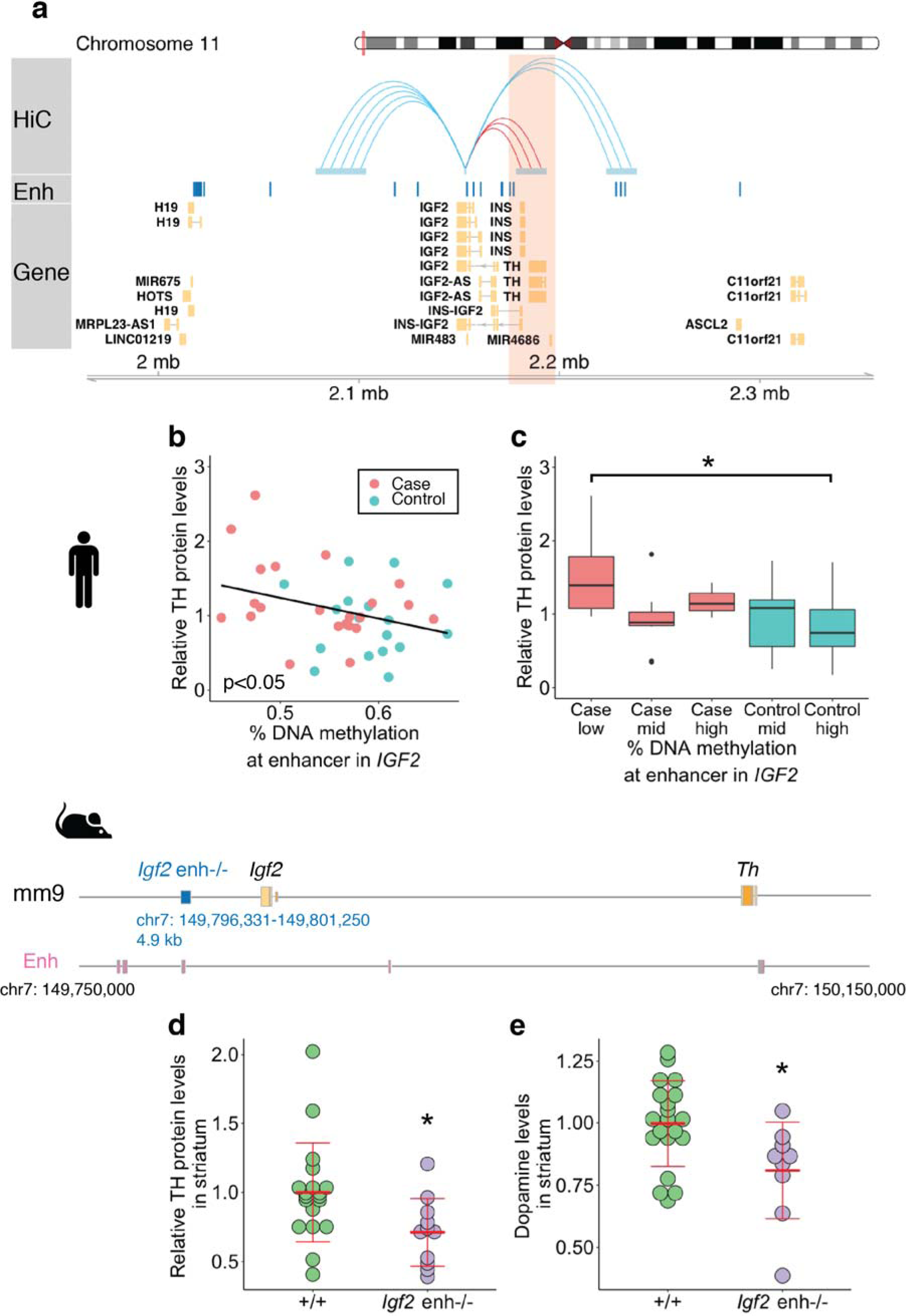
The enhancer at *IGF2* found differentially methylated in major psychosis regulates the dopamine synthesis enzyme tyrosine hydroxylase (*TH*). (**a**) Interaction of the *IGF2* enhancer with the *TH* gene. Chromatin interactions of enhancer at the *IGF2* locus that is differentially methylated in major psychosis were identified in the prefrontal cortex using Hi-C analysis reported on the 3D Interaction Database (https://www.kobic.kr/3div/); interactions within ± 100 kb are depicted in the figure. Chromatin states in the adult brain prefrontal cortex tissue are shown (Enh: enhancer; data from NIH Epigenomics Roadmap ChromHMM 18-state model). (**b**) Reduced DNA methylation at the *IGF2* enhancer in major psychosis correlates to an increase in TH protein levels (*p*<0.05 by linear regression). Immunoblotting was used to measure TH in control (red dots, n = 17) and major psychosis (blue dots, n = 22) prefrontal cortex. TH protein levels are relative to control proteins (NeuN, INA, actin). (**c**) DNA methylation status at *IGF2* enhancer results in TH protein levels differences between major psychosis cases and controls (Same data as (b)). Low, mid, or high DNA methylation (<50%, 50-60%, and >60%, respectively) at the *IGF2* enhancer in cases and controls are shown (left to right n = 8, 11, 3, 9, and 8). Main effect of DNA methylation by two-way ANOVA *F*_(2, 35)_=3.5, *p*<0.05; \**p*=0.05 by Tukey post-hoc test. (**d** and **e**) Effect of *Igf2* enhancer knockout in mice. Schema shows deletion of the 4.9 kb *Igf2* enhancer alongside mouse forebrain enhancers (ENCODE). **(d)** Relative TH protein levels in striatum of adult wild-type (+/+; n = 19) and *Igf2*enh-/-(n = 11) mice (NeuN, actin control). **(e)** Dopamine levels in striatum of adult wild-type (n = 20) and *Igf2*enh-/-(n = 9) mice measured by HPLC. \**p*<0.05 by one-way ANOVA.

In sum, we have identified an increase in permissive epigenetic marks at an enhancer regulating the *TH* gene in neurons of patients with major psychosis. Enhancer-mediated upregulation of TH causing higher striatal dopamine synthesis would augment the risk for psychosis^8^. Interestingly, in patients, the progressive loss of prefrontal cortex volume closely parallels the development of psychosis^10, 11^. Imaging studies of at-risk individuals show greater prefrontal cortical volume loss in individuals that transition to psychosis compared to those remaining healthy^11^. The severity of psychotic symptoms is also associated with structural alterations in the cortex^12^. In the brain, *IGF2* promotes synapse development, spine maturation, and memory formation^13-16^, signifying that normal *IGF2* activation is required for healthy neuronal architecture. Recently, *IGF2* was found to be the top downregulated gene in the schizophrenia prefrontal cortex in the large CommonMind consortium RNA-sequencing study^17^. Loss of DNA modification at the *IGF2* locus has been associated with decreased *IGF2* mRNA levels in early development^18^, and risk factors for schizophrenia; prenatal exposure to famine^19^ and reduced brain weight^20^. Similarly, our transcriptome analysis found a downregulation of genes affecting synaptic transmission and interacting with *IGF2*. Therefore, in neurons of major psychosis patients, epigenetic changes that facilitate the recruitment of the enhancer at *IGF2* for activation of the *TH* gene, may impede *IGF2* regulation in tandem. We propose a model in which improper epigenetic control of an *IGF2* enhancer simultaneously contributes to dopamine-mediated psychotic symptoms and synaptic structural deficits in major psychosis.

## Acknowledgements

S.P. and V.L. are supported by the Brain & Behavior Research Foundation (529941 to S.P.; 23482 to V.L.). V.L. is also supported by grants from the Alzheimer’s Society of Canada (16 15) and Scottish Rite Charitable Foundation of Canada (15110). We thank John Murdoch for assistance with targeted bisulfite sequencing library preparation. We thank Van Andel Research Institute core services including the pathology and biorepository, genomics, and bioinformatics and biostatistics. We also thank Therese Murphy, Jonathan Mill and Sarah Gagliano for feedback on parts of this work, and Gary D. Bader for his support.

We gratefully acknowledge the tissue banks, donors and families from whom these samples were obtained. Tissue samples were obtained from the following tissue banks through the NIH NeuroBioBank at: the Harvard Brain Tissue Resource Center (supported in part by PHS contract, HHSN-271-2013-00030C); the Human Brain and Spinal Fluid Resource Center (VA West Los Angeles Healthcare Center, Los Angeles CA 90073 which is sponsored by NINDS/NIMH, National Multiple Sclerosis Society, and the Department of Veterans Affairs); the University of Miami Brain Endowment Bank; and the University of Pittsburgh Brain Tissue Donation program.

Histone peaks from CNON cells were generated as part of the PsychENCODE Consortium, supported by: U01MH103339, U01MH103365, U01MH103392, U01MH103340, U01MH103346, R01MH105472, R01MH094714, R01MH105898, R21MH102791, R21MH105881, R21MH103877, and P50MH106934 awarded to:

Schahram Akbarian (Icahn School of Medicine at Mount Sinai), Gregory Crawford (Duke), Stella Dracheva (Icahn School of Medicine at Mount Sinai), Peggy Farnham (USC), Mark Gerstein (Yale), Daniel Geschwind (UCLA), Thomas M. Hyde (LIBD), Andrew Jaffe (LIBD), James A. Knowles (USC), Chunyu Liu (UIC), Dalila Pinto (Icahn School of Medicine at Mount Sinai), Nenad Sestan (Yale), Pamela Sklar (Icahn School of Medicine at Mount Sinai), Matthew State (UCSF), Patrick Sullivan (UNC), Flora Vaccarino (Yale), Sherman Weissman (Yale), Kevin White (UChicago) and Peter Zandi (JHU)."

## Software versions, URLs and references

Software URLs and references are provided in a single table at the end of the Online Methods section.

## Accessions code/data availability

Software used to produce these results is available at https://github.com/shraddhapai/EpigeneticsPsychosis and will be made public upon publication. Methylation arrays generated in this project have been deposited at GEO under accession number GSE112179 and GSE112525.

## Author Contributions

The study was designed and coordinated by S.P. and V.L. V. L. and P.J. contributed to the sorting of neuronal nuclei and DNA extraction for epigenetic and genetic experiments. S.P. performed the computational analysis for the epigenetic, genetic and transcriptomic datasets. L.M. processed the transcriptomic dataset. B.A.K. performed immunoblotting and HPLC experiments. J.L. and P.S. generated and managed the transgenic mice with the *Igf2* enhancer deletion. P.L. analyzed the chromatin conformation data and contributed to the DNA modification analysis. A.P. helped oversee the project. The manuscript was written by S.P. and V.L., and commented on by all authors.

## Competing financial interests

The authors declare no competing financial interests.

## Online Methods

### Human brain samples

Post-mortem brain samples were obtained through the NIH NeuroBioBank at the University of Pittsburgh; the Harvard Brain Tissue Resource Center; the Human Brain and Spinal Fluid Resource Center at Sepulveda; and the University of Miami Brain Endowment Bank. We obtained sample information on demographic factors (age, sex), clinical variables (cause of death, medications at time of death, duration of antipsychotic use, smoking status, brain weight), and tissue quality (post-mortem interval, tissue quality/RIN score). Patient data is provided in Supplementary Table 1. Approximately equal numbers of males and females were used in control and case groups. Our analyses controlled for sample age, sex, post-mortem interval, ethnicity (and batch effect in array-based studies), and the influence of the clinical covariates was examined in our data. The study protocol was approved by the institutional review board at the Centre for Addiction and Mental Health and the Van Andel Research Institute (IRB #15025).

### Isolation of neuronal nuclei using a flow cytometry

Neuronal nuclei were separated using a flow cytometry-based approach, similar to as previously described^1, 2^. Human brain tissue (250 mg) for each sample was minced in 2 mL PBSTA (0.3 M sucrose, 1X phosphate buffered saline (PBS), 0.1% Triton X-100). Samples were then homogenized in PreCellys CKMix tubes with a Minilys (Bertin Instruments) set at 3,000 rpm for three 5 sec intervals, 5 min on ice between intervals. Samples homogenates were filtered through Miracloth (EMD Millipore), followed by a rinse with an additional 2 mL of PBSTA. Samples were then placed on a sucrose cushion (1.4 M sucrose) and nuclei were pelleted by centrifugation at 4,000 × g for 30 min 4°C using a swinging bucket rotor. For each sample, the supernatant was removed and the pellet was incubated in 700 μl of 1X PBS on ice for 20 min. The nuclei were then gently resuspended and blocking mix (100 μl of 1X PBS with 0.5% BSA (Thermo Fisher Scientific) and 10% normal goat serum (Gibco) was added to each sample. NeuN-488 (1:500; Abcam) was added and samples were incubated 45 min at 4°C with gentle mixing. Immediately prior to flow cytometry sorting, nuclei were stained with 7-AAD (Thermo Fisher Scientific) and passed through a 30 μM filter (SystemX). Nuclei positive for 7-AAD and either NeuN+ (neuronal) or NeuN-(non-neuronal) were sorted using an Influx (BD Biosciences) at the Faculty of Medicine Flow Cytometry Facility at the University of Toronto (Toronto, ON, Canada). Approximately 1 million NeuN+ nuclei were sorted for each sample. Immediately, after sorting nuclei were placed on ice and then precipitated by raising the volume to 10 mL with 1X PBS and adding 2 mL 1.8 M sucrose, 50 μl 1M CaCl_2_ and 30 μl Mg(Ace)_2_ and centrifugation at 1,786 × g for 15 min at 4°C. The supernatant was removed from NeuN+ and NeuN-samples and pellets were stored at -80°C. Genomic DNA from each NeuN+ and NeuN-fraction of each sample was isolated using standard phenol-chloroform extraction methods.

### Genome-wide DNA methylation analysis

Whole-genome DNA methylation profiling for each sample was performed on Illumina MethylationEPIC BeadChip microarrays at The Centre for Applied Genomics (Toronto, Canada). Bisulfite converted DNA samples (n=104) were randomized across arrays (8 samples/array). Data generated from the microarrays were preprocessed with Minfi v1.19.12. Normalization was performed with noob^3^, followed by quantile normalization. All samples had sex matching that predicted from the methylome. Probes that overlapping SNPs (minor allele frequency > 5%) on the CpG or single-base extension were excluded (11,812 probes), as were probes known to be cross-reactive^4^ (42,558 probes) and those that failed detectability (P>0.01) in >20% samples (1,170 probes). After processing, 812,663 probes were left. Based on principal component analysis (PCA) co-clustering, one sample, despite being labeled NeuN+, clustered with NeuN-samples and was excluded from downstream analyses. Surrogate variable analysis identified no such variables; for this the null model used age and sex as covariates, while the full model included diagnosis.

The top 50% probes with highest variance were used to identify differentially methylated probes (406,332 probes). For each probe, a linear model was fit using limma^5^; in addition to diagnosis, age, sex, post-mortem interval and first two principal components of genetic ancestry were used as covariates. Ethnicity covariates were computed by PCA within plink^6, 7^. As the design included several technical replicates, these were modeled as blocking factors. Variance shrinkage was applied (limma in R). Benjamini Hochberg FDR correction was used to correct nominal p-values. Analysis was performed in R (3.3.1). To identify differentially methylated regions (DMRs) in our data, we used the Python module Comb-p^8^ to group spatially correlated differentially methylated probes (seed p-value of 0.01 to start a region, at a maximum distance of 500bp in each brain tissue). The input data was a sorted BED with a column for p-value from single-probe tests. DMR p-values were corrected for multiple testing using Šidák correction.

### Targeted bisulfite sequencing at IGF2 locus

DNA methylation at the *IGF2* and surrounding genomic area (161 kb) was captured using the SeqCap Epi Enrichment System (Roche). Biotinylated long oligonucleotide probes targeting 450 sites at the extended *IGF2* locus (unique, non-repetitive genome) were custom designed by Roche NimbleGen. Library preparation was performed following manufacturer instructions. Briefly, gDNA (500 ng) of each sample (n= 4 cases and 3 controls, in addition 4 technical replicate samples) were fragmented (~ 200 bp), end repaired and ligated to barcoded adapters using the KAPA Library Preparation kit (Kapa Biosystems) and SeqCap Adapter Kit A and B (Roche). Bisulfite conversion of the adapter ligated DNA, followed by column purification, was performed with the EZ DNA Methylation Lightning kit (Zymo). The bisulfite converted DNA for each sample was then amplified by ligation mediated PCR (95°C 2 min, 10 cycles of [98°C 30 sec, 60°C 30 sec, 72°C 4 min], 72°C 10 min, 4°C hold) followed by purification with Agencourt AMPure XP beads (Beckman Coulter). Sample quality was verified on Bioanalyzer (Agilent) and quantity was determined with a NanoDrop spectrophotometer (Thermo Fisher Scientific). Equimolar amounts of each sample were then combined into a single pool. The *IGF2* target region was captured by hybridizing the amplified bisulfite converted DNA pool (1 μg) to the probe library (Roche), as directed by manufacturer. Enrichment and recovery of captured bisulfite-converted DNA was completed by binding to magnetic beads and subsequent wash steps using the SeqCap Pure Capture Bead kit and the SeqCap Hybridization and Wash kit (Roche). The captured DNA was then amplified by ligation mediated PCR (98°C 45 sec, 11 cycles of [98°C 15 sec, 60°C 30 sec, 72°C 30 sec], 72°C 1 min) followed by purification with Agencourt AMPure XP beads (Beckman Coulter). Library quality and quantity was assessed using a combination of Agilent DNA High Sensitivity chip on a Bioanalyzer (Agilent Technologies), Qubit dsDNA HS Assay kit on a Qubit 3.0 fluorometer (Thermo Fisher Scientific), and Kapa Illumina Library Quantification qPCR assays (Kapa Biosystems). DNA sequencing was performed on an Illumina HiSeq 2500 on Rapid Run mode with all samples run on both sequencing lanes.

Data were processed using the pipeline recommended by the manufacturer^9^. Trimmomatic^10^ was used to trim read adapter sequences and BSMAP^11^ was used to align reads to the GRCh38/hg38 genome build. The genome index consisted of reference chromosome sequences and the lambda phage genome (https://www.ncbi.nlm.nih.gov/nuccore/215104). Following alignment, reads were pooled across the two lanes. The merged set of reads was separated into those aligning to top and bottom strand, duplicates were removed with Picard, and then matching read pairs were merged. Samtools^12^ was used to exclude reads that were not properly paired or were unmapped. Bamutils were used to clip overhanging reads that distort methylation estimates. Methratio.py in BSMAP computed the percent methylation at the base level.

### meQTL analysis

SNPs in each sample (n=104) were determined using the Infinium PsychArray-24 processed by The Centre for Applied Genomics (Toronto, Canada). Samples were randomized across SNP arrays. LiftOver was used to convert genotypes to the GRCh37/hg19 build. Quality control was performed as described^13^. SNPs with a minor allele frequency < 0.05, those with HWE *p* < 10^−6^ and those missing in >1% samples were excluded. Where pairs of individuals had relatedness (Identity By State; IBS) > 0.185, one was excluded. Samples with < 90% SNPs genotyped and those with outlier heterozygosity were excluded. 96 samples passed these filters and were used for meQTL analysis. PCA for EWAS and meQTL analysis was extracted using plink on study samples. Continental genetic ancestry was ascertained by MDS using HapMap3 as a reference population^14^. For European-specific EWAS, Europeans were defined as individuals with MDS 1 and 2 lying within 3 standard deviations of the mean defined by the CEU population in the HapMap3 reference panel. For meQTL analysis and allele-specific expression analysis, we first imputed genotype data using Check-bim and the Michigan Imputation Server^15^ (Eagle v2.3^16^; 1000G^17^ Phase 3 v5; Population:ALL). SNPs with INFO score > 0.7 were retained.

For cis e-QTL inference, SNPs within +/-500 kb of CpG probes in differentially-methylated regions were tested. For trans e-QTL inference, SNPs from schizophrenia GWAS study^18^ (SNPs with nominal *p* < 10^−9^), and SCZ “credible” SNPs were included^18, 19^. Only SNPs with ≥10 individuals per genotype were tested. A linear regression was used to assess the effect of genotype on methylation, with sex, diagnosis, age, and the first two genetic principal components as covariates. Methylation for technical replicates was averaged, and where technical replicates existed for genotype, the first replicate was used. Benjamini Hochberg correction was applied for multiple testing.

### Gene expression profiling by RNA-seq

Brain tissue samples (n=34 human samples) were lysed using QIAzol Lysis Reagent (Qiagen) and homogenized with a TissueLyser (Qiagen). Total RNA from each sample was isolated using the RNeasy Plus Universal Mini kit (Qiagen) according to manufacturer’s instructions and included an enzymatic DNase (Qiagen) digestion step. RNA quality was measured on a 2100 Bioanalyzer (Agilent) and quantity was determined with a Nanodrop 2000 spectrophotometer (Thermo Fisher Scientific). RNA samples had a RIN quality score >7 and proceeded to RNA-seq library preparation (RIN between 7.1 to 9.4 for all samples). Libraries were prepared by the Van Andel Genomics Core from 300 ng of total RNA using the KAPA RNA HyperPrep Kit with RiboseErase (v1.16) (Kapa Biosystems). RNA was sheared to 300-400 bp. Prior to PCR amplification, cDNA fragments were ligated to Bio Scientific NEXTflex Adapters (Bioo Scientific). Quality and quantity of the finished libraries were assessed using a combination of Agilent DNA High Sensitivity chip (Agilent Technologies, Inc.), QuantiFluor^®^ dsDNA System (Promega Corp.), and Kapa Illumina Library Quantification qPCR assays (Kapa Biosystems). Individually indexed libraries were pooled, and 75 bp paired-end sequencing was performed on an Illumina NextSeq 500 sequencer, with all libraries run across 3 flowcells. Base calling was done by Illumina NextSeq Control Software (NCS) v2.0 and output of NCS was demultiplexed and converted to FastQ format with Illumina Bcl2fastq v1.9.0.

Trimgalore (v0.11.5) was used for adapter removal prior to genome alignment. STAR^20^ (v2.3.5a) index was generated using Ensemble GRCh38 p10 primary assembly genome and the Gencode v26 primary assembly annotation. Read alignment was performed using a STAR two-pass mode. Gene counts matrix was imported into R (3.4.1) and low expressed genes (counts per million (CPM) < 1 in all samples) were removed prior to differential expression in EdgeR. Gene counts were normalized using the trimmed mean of M-values, fitted in a generalized linear model and differentially tested using a likelihood ratio test. The generalized linear model contained covariates age, sex, post mortem interval and neuronal cell composition. Cell-type compositions for each sample was accessed using CIBERSORT^21^ on normalized sample counts against cell-type specific markers (see below), identifying the proportion of neurons in each samples. Benjamini Hochberg correction was used to adjust for multiple testing.

Our RNA-seq analysis corrected for the proportion of neuronal cells in each sample. Neuronal cell proportions were determined by CIBERSORT^21^ (http://cibersort.stanford.edu), which involved a gene signature matrix derived from single cell RNA-seq measures in adult human brain cells (signature matrix^22^; source^23^). Because major psychosis is characterized by a loss of synaptic density, we excluded genes encoding synaptic proteins (Genes2Cognition database^24^; lists L00000009, L00000016, L00000012) from the gene signatures. 135 synapse-associated genes were excluded, leaving 768 genes in the deconvolution analysis. CIBERSORT was run (100 permutations), and the inferred proportion of neurons was used as a covariate for differential expression.

For allele specific expression (ASE) analysis, we used Samtools and Picard to prepare RNAseq BAM files for ASE read counting. ASEReadCounter was used to count allele-specific transcript counts. Allelic imbalance was defined as |0.5-REF_COUNT/(REF_COUNT+ALT_COUNT)|, and only sites with one or more heterozygous alleles were included.

### Pathway enrichment analysis

Pathways affected by the DNA methylation and transcriptomic changes in major psychosis were determined. For DNA methylation data, probes were mapped to genes if they overlapped with [TSS-1 kb, TES]. Gencode^25^ v27 (liftOver to GRCh37) were used for gene extents. The pathway file was the same as that used for RNA pathway analysis; pathways with 10-500 genes were included. For each pathway, a hypergeometric test was performed comparing the proportion of foreground probes (*p* < 0.05 from DMR analysis) to background probes (all probes tested for DMRs). Benjamini-Hochberg correction was performed to adjust for multiple testing.

For gene expression data, gene set enrichment analysis (GSEA^26^) was performed (1000 permutations) to identify the potentially dysregulated pathways. Pathway definitions were aggregated from HumanCyc^27^, IOB’s NetPath^28^, Reactome^29, 30^, NCI Curated Pathways^31^, mSigDB^26^, Panther^32^, and Gene Ontology^33, 34^. Genesets with 10-200 genes were included (5,672 pathways).

### Igf2 enhancer deletion in mice

A 4.9 kb-long DNA fragment (chr7: 149,796,331-149,801,250 in mm9) was deleted from the intergenic region of *H19* and *Igf2* by classical ES cell gene targeting in the mouse in the 129S1 genetic background. One *loxP* site remained at the site of the deletion mutation after the excision of the Pgkneo positive selection cassette by crossing the targeted mutant male mouse to an Hprt-CRE transgenic female^35^. Three oligonucleotide primers, IGKOCrerecU: CGGAATGTTTGTGTGGAGAGCA; IGKOwtU: TAGGGGTCCTGAAGACGTCAG; and IGKOCreWTL: TTGGTGTAGCACCCTGTAACCC are combined in one PCR reaction to distinguish the mutant from the wild type allele, as visualized by a 450 bp or a 350 bp long PCR product, respectively.

Mice were bred and housed in ventilated polycarbonate cages, and given *ad libitum* sterile food (LabDiet 5021) and water. Adult mice were housed by sex in groups of 2-5 littermates. The vivarium was maintained under controlled temperature (21°C±1°C) and humidity (50-60%), with a 12-h diurnal cycle (lights on: 0700-1900). Approximately equal numbers of male and female were tested, and no sex differences were detected (in all Western blotting and HPLC experiments). No animals were excluded from the study. Wild-type (+/+) and homozygote knock-out (*Igf2* enh-/-) adult mice (~2.5 months old) were tested, and sample sizes were comparable to other studies of *Igf2* mutant mice^36, 37^. All animal procedures were approved by the Institutional Animal Care Committee of the Van Andel Research Institute and complied with the requirements of the Institutional Animal Care and Use Committee (AUP # PIL-17-10-010).

### Immunoblotting

All tissue preparation procedures were performed on ice. Frozen tissue samples weighing ~20 mg were sonicated in 500 µl of RIPA buffer (10 mM Tris-HCl pH 8, 1% Triton X-100, 0.1% sodium deoxycholate, 0.1% SDS, 140 mM NaCl, protease inhibitor cocktail from Roche, and 1 mM EDTA) and incubated for 1 h on ice with mixing. The samples were then centrifuged at 22,000 × g for 30 min. Protein content of the supernatant was determined using BCA assay (Thermo Fisher Scientific) and then diluted in SDS-PAGE sample buffer (Biorad) to yield 20 µg protein per lane. Samples were separated on 4-20% SDS-PAGE gels (Thermo Fisher Scientific) and blotted onto 0.22 µm PVDF membranes (Thermo Fisher Scientific) for 2 h at a constant 20 V using xCell II blot module (Thermo Fisher Scientific). Membranes were blocked in TBST (50 mM Tris-HCl pH 7.6, 150 mM NaCl, and 1% tween-20) containing 5% non-fat dry milk (Bio-Rad) for 1 h at room temperature. Membranes were then incubated with blocking buffer containing primary antibodies: tyrosine hydroxylase (TH, PelleFreez Biologicals, AR), NeuN (Cell Signaling), internexin neuronal intermediate filament protein (INA, Sigma), and actin (Millipore) diluted 1:1000 overnight at 4°C. Membranes were washed three times for five minutes with TBST and probed with the appropriate HRP-conjugated anti-igg antibodies according to manufacturers recommended protocol. Blots were then washed three times for 5 min with TBST and imaged using west pico ECL reagent (Thermo Fisher Scientific).

### Quantification of dopamine levels by high performance liquid chromatography (HPLC)

All tissue preparation procedures were performed on ice. Frozen tissue samples weighing between 5-20 mg were sonicated in 100-300 µl of 0.2 M perchloric acid (Sigma). The sample was centrifuged at 22,000 x g for 30 min and the resulting supernatant was filtered using 0.22 µM cellulose acetate filter (Costar). The filtered supernatant was separated using the HTEC-500 HPLC system (Eicom) with the SC-30DS reverse phase separation column (Eicom) and electrochemical detector. Samples were separated in mobile phase consisting of 0.1 M citrate acetate pH 3.5, 20% methanol, 220 g/L sodium octane sulfonate, and 5 mg/L EDTA-Na. The samples were then compared to known standards of dopamine (Sigma), homovinillic acid (HVA), and 3,4-Dihydroxyphenylacetic acid (DOPAC). The pellet was dissolved in 0.5 mL of 1 M NaOH for 10 min at 90°C, and the resulting protein concentration determined by BCA assay. The final values were calculated as ng analyte per µg protein.

### Lifestyle factors

FDA-approved antipsychotics were collected from the literature (https://www.fda.gov/Drugs/DrugSafety/ucm243903.htm,^38^), including generic and brand names. Patient medication information was computationally searched for keyword matches from this list to identify drugs used by individuals; the mood stabilizers lithium and valproic acid were added to this list. Where no match was found, antipsychotic status was set to “none”. Analysis divided patients into those who ever had antipsychotic use and those who did not. Smoking status was binarized so that any record of lifetime smoking resulted in a categorization of the sample as a smoker. Individuals with missing information were put into a separate category.

#### Olfactory neuroepithelial Chip-Seq

Peak calls for histone marks in DNA isolated from olfactory neuroepithelial cells (CNON) in controls and individuals diagnosed with schizophrenia were downloaded from the PsychENCODE^39^ knowledge portal (https://www.synapse.org/#!Synapse:syn4590897). The average of peak intensities in biological replicates was computed using a 250bp tiling window.

### Software used for analysis

| Name | Ver. | URL | Ref |
| --- | --- | --- | --- |
| BSMap | 2.74 | https://code.google.com/archive/p/bsmap/downloads | 11 |
| Picard | 2.9.4-SNAPSHOT | https://github.com/broadinstitute/picard<br>http://picard.sourceforge.net |  |
| Bedtools | 2.26 | https://github.com/arq5x/bedtools2 | 40 |
| Samtools | 1.5 | http://www.htslib.org/ | 12 |
| FastQC | v0.11.5 | https://www.bioinformatics.babraham.ac.uk/projects/fastqc/ |  |
| Trimmomatic | 0.36 | http://www.usadellab.org/cms/?page=trimmomatic | 10 |
| Tabix (HTSUtils) | 1.5 | http://www.htslib.org/ | 41 |
| edgeR | 3.18.1 | http://bioconductor.org/packages/release/bioc/html/edgeR.html | 42 |
| STAR | 2.5.3a | https://github.com/alexdobin/STAR | 20 |
| Minfi | 1.22.1 | http://bioconductor.org/packages/release/bioc/html/minfi.html | 43 |
| Cytoscape | 3.5.1 | http://cytoscape.org/ | 44 |
| GSEA | 3.0 | http://software.broadinstitute.org/gsea/index.jsp | 26 |
| EnrichmentMap | 3.1.0RC4 | http://apps.cytoscape.org/apps/enrichmentmap | 45 |
| AutoAnnotate | 1.2 | http://apps.cytoscape.org/apps/autoannotate | 46 |
| CIBERSORT |  | http://cibersort.stanford.edu | 21 |
| Plink | 1.90b4.9 | https://www.cog-genomics.org/plink/1.9/ | 7 |
| Michigan Imputation Server |  | https://imputationserver.sph.umich.edu/index.html | 15 |
| Eagle | 2.3 | (part of Michigan Imputation Server) | 16 |
| Check-Bim |  | http://www.well.ox.ac.uk/~wrayner/tools/#Checking |  |
| Pathway databases |  | http://humancyc.org<br>http://www.netpath.org<br>http://www.reactome.org<br>http://software.broadinstitute.org/gsea/msigdb/<br>http://pantherdb.org/<br>http://download.baderlab.org/EM_Genesets/October_01_2017/Human/symbol/Human_GOBP_AllPathways_no_GO_iea_October_01_2017_symbol.gmt | 26-32, 45 |

## Supplementary Figures

**Supplementary Figure 1.**
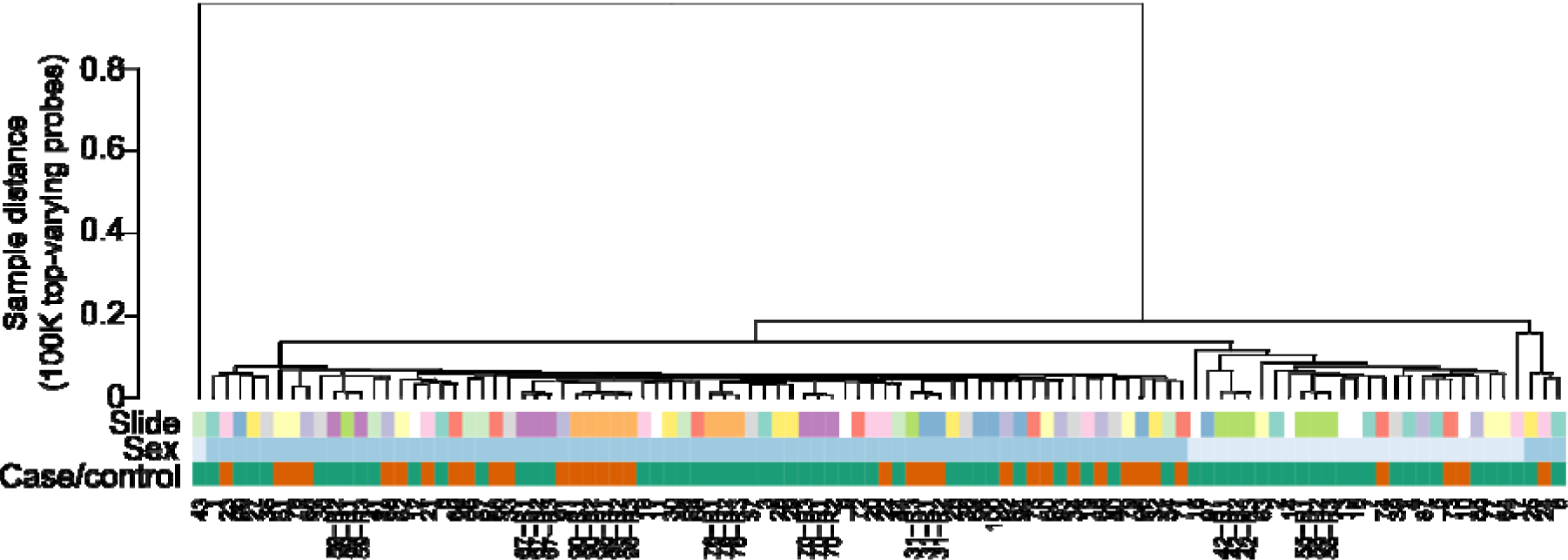
Hierarchical clustering of MethylationEPIC array samples using 100K probes with highest variance. Samples are coded by microarray slide, sex, and case/control status. Sample suffixed with –R1,2,3 are technical replicates. All samples are NeuN+ except for one NeuN-sample at the left branch of plot.

**Supplementary Figure 2.**
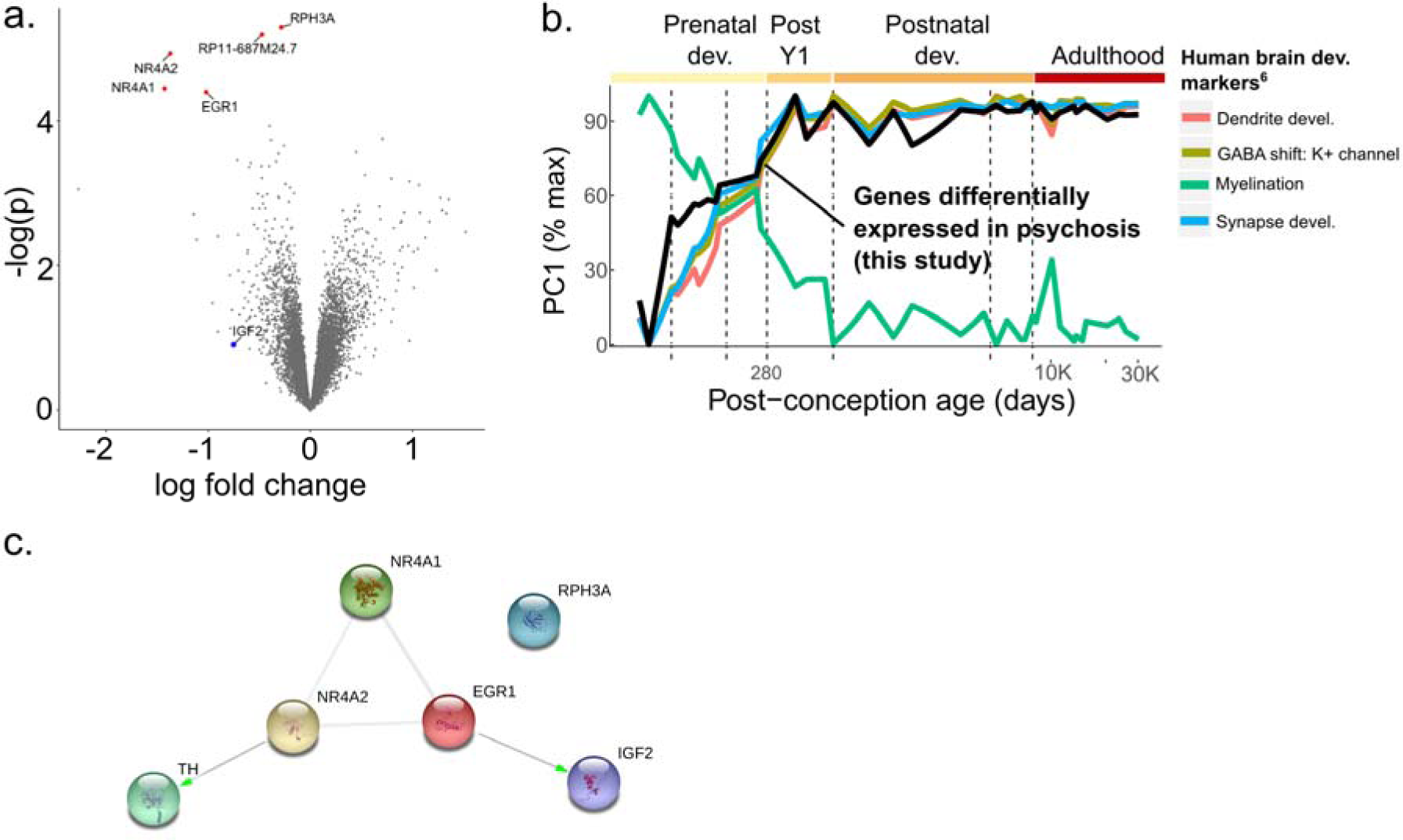
Transcriptional changes in major psychosis identified by RNA-sequencing. (**a**) Volcano plot of differential expression analysis between major psychosis cases (n=17) and controls (n=17), after controlling for age, sex, post-mortem interval and neuronal cell percentage. Dots in red have *p* < 10^−4^; *IGF2* is highlighted in blue. (**b**) Transcript level changes in human frontal cortex across the life-cycle for genes that were differentially expressed in major psychosis. Data from BrainSpan, which included the human frontal cortex and areas of the ganglionic eminence. Genes associated with a developmental process were obtained from BrainSpan (Table S13 of that work). Each trendline shows the first principal component of the gene-set, scaled from 0-100 across the lifespan. The black trendline represents the genes differentially expressed in the current study (p<0.05, n=854 genes) (**c**) Top differentially expressed genes in major psychosis have known protein-protein interactions with *IGF2* and *TH* (image from STRING database).

**Supplementary Figure 3.**
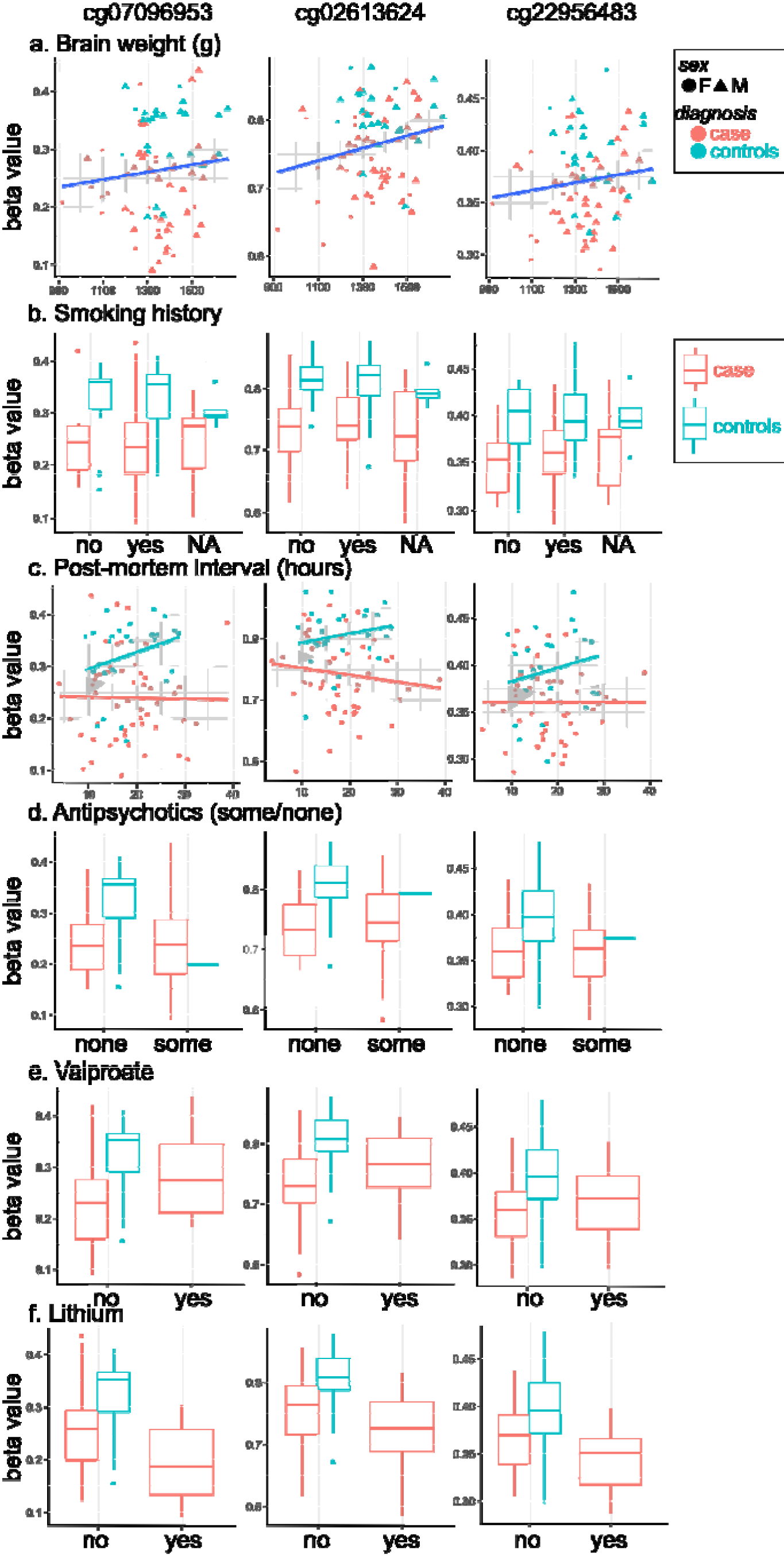
*IGF2* enhancer methylation considering lifestyle factors. Each column shows data for one of the three top probes at the *IGF2* locus differentially methylated in major psychosis. Rows show the effect of brain weight (**a**), smoking (**b**), post-mortem interval (**c**), antipsychotics use (**d**), valproate use (**e**), and lithium use (**f**). Y-axis shows probe-level beta-values. Controls are shown in teal and cases are shown in pink.

**Supplementary Figure 4.**
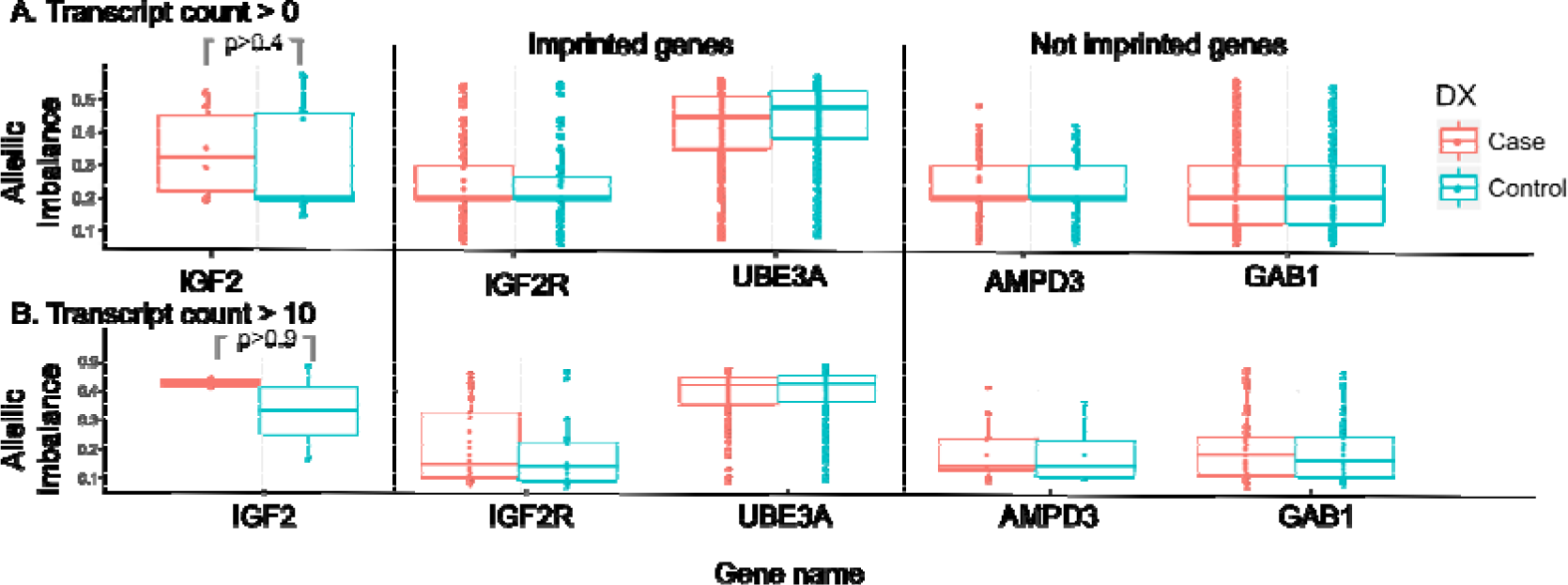
Allele-specific differences in *IGF2* expression in the frontal cortex are similar between major psychosis cases and controls. Allelic imbalances in major psychosis cases (n=17) and controls (n=17). Shown are results for the *IGF2* gene, and for comparison, two other known imprinted genes and two genes that are not imprinted. Each boxplot shows data for individual CpGs and samples (unaggregated), such that each dot represents the measure at one site in one sample. (a) Sites not filtered for transcript count. (b) Sites filtered to those with >=10 reads.

**Supplementary Figure 5.**
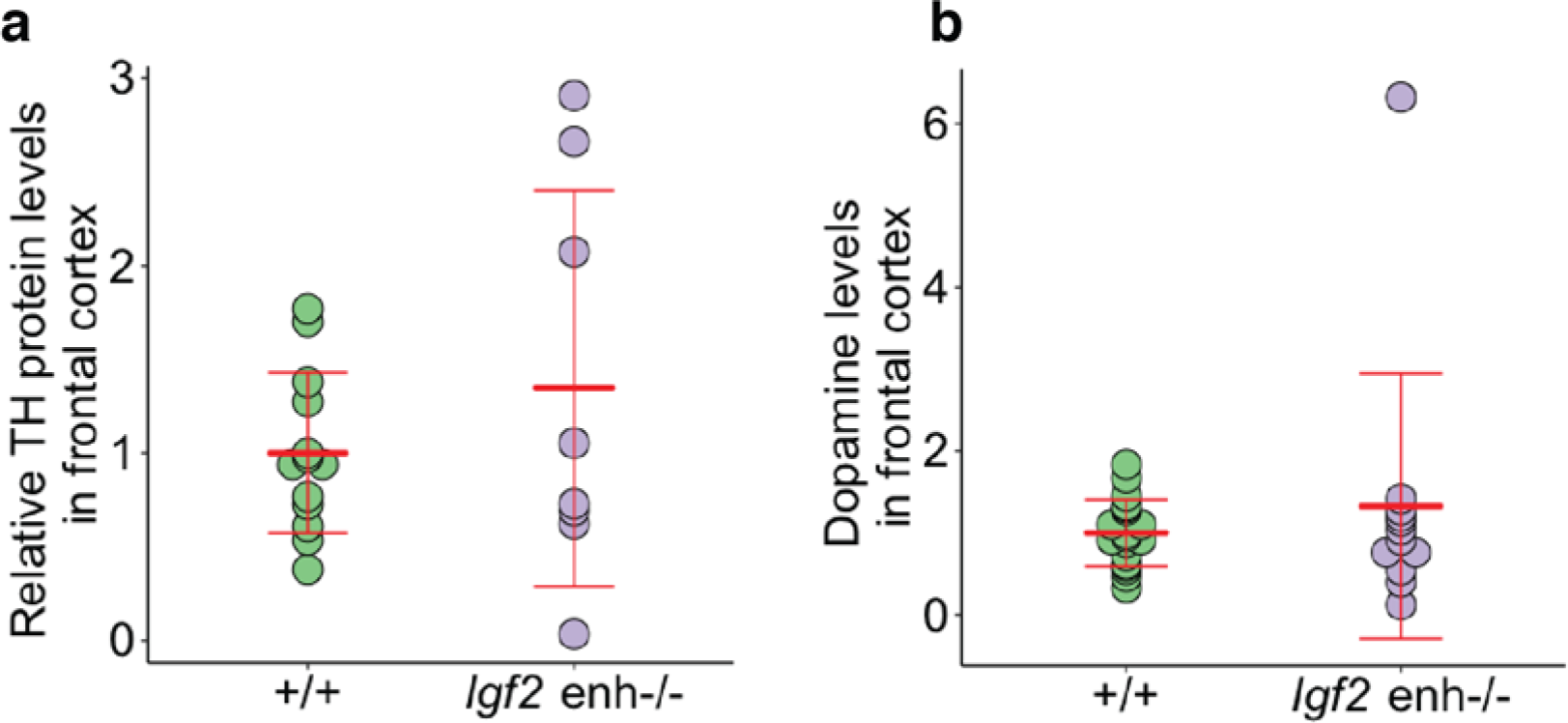
TH protein and dopamine levels are not altered in the frontal cortex of mice with an *Igf2* enhancer deletion. (**A**) Immunoblotting was used to measure TH protein levels in the frontal cortex of adult *Igf2*enh-/-(n = 13) and wild-type (+/+; n = 8) mice. TH protein levels are relative to control proteins (NeuN, actin). (**B**) Dopamine levels in the frontal cortex of adult wild-type (n = 19) and *Igf2*enh-/-(n = 12) mice were measured by HPLC.

## References

1. Hannon, E., et al. Methylation QTLs in the developing brain and their enrichment in schizophrenia risk loci. Nature neuroscience 19, 48–54 (2016).

2. Jaffe, A.E., et al. Mapping DNA methylation across development, genotype and schizophrenia in the human frontal cortex. Nature neuroscience 19, 40–47 (2016).

3. Ludwig, B. & Dwivedi, Y. Dissecting bipolar disorder complexity through epigenomic approach. Molecular psychiatry 21, 1490–1498 (2016).

4. Wockner, L.F., et al. Genome-wide DNA methylation analysis of human brain tissue from schizophrenia patients. Transl Psychiatry 4, e339 (2014).

5. Montano, C., et al. Association of DNA Methylation Differences With Schizophrenia in an Epigenome-Wide Association Study. JAMA Psychiatry 73, 506–514 (2016).

6. Chen, C., et al. Correlation between DNA methylation and gene expression in the brains of patients with bipolar disorder and schizophrenia. Bipolar Disord 16, 790–799 (2014).

7. Ribeiro, P.F., et al. The human cerebral cortex is neither one nor many: neuronal distribution reveals two quantitatively different zones in the gray matter, three in the white matter, and explains local variations in cortical folding. Front Neuroanat 7, 28 (2013).

8. Howes, O.D., et al. The nature of dopamine dysfunction in schizophrenia and what this means for treatment. Arch Gen Psychiatry 69, 776–786 (2012).

9. Ashok, A.H., et al. The dopamine hypothesis of bipolar affective disorder: the state of the art and implications for treatment. Molecular psychiatry 22, 666–679 (2017).

10. Olabi, B., et al. Are there progressive brain changes in schizophrenia? A meta-analysis of structural magnetic resonance imaging studies. Biological psychiatry 70, 88–96 (2011).

11. Sun, D., et al. Progressive brain structural changes mapped as psychosis develops in ’at risk’ individuals. Schizophr Res 108, 85–92 (2009).

12. Satterthwaite, T.D., et al. Structural Brain Abnormalities in Youth With Psychosis Spectrum Symptoms. JAMA Psychiatry 73, 515–524 (2016).

13. Schmeisser, M.J., et al. IkappaB kinase/nuclear factor kappaB-dependent insulin-like growth factor 2 (Igf2) expression regulates synapse formation and spine maturation via Igf2 receptor signaling. The Journal of neuroscience : the official journal of the Society for Neuroscience 32, 5688–5703 (2012).

14. Ferron, S.R., et al. Differential genomic imprinting regulates paracrine and autocrine roles of IGF2 in mouse adult neurogenesis. Nature communications 6, 8265 (2015).

15. Terauchi, A., Johnson-Venkatesh, E.M., Bullock, B., Lehtinen, M.K. & Umemori, H. Retrograde fibroblast growth factor 22 (FGF22) signaling regulates insulin-like growth factor 2 (IGF2) expression for activity-dependent synapse stabilization in the mammalian brain. Elife 5 (2016).

16. Chen, D.Y., et al. A critical role for IGF-II in memory consolidation and enhancement. Nature 469, 491–497 (2011).

17. Fromer, M., et al. Gene expression elucidates functional impact of polygenic risk for schizophrenia. Nature neuroscience 19, 1442–1453 (2016).

18. Murrell, A., et al. An intragenic methylated region in the imprinted Igf2 gene augments transcription. EMBO reports 2, 1101–1106 (2001).

19. Tobi, E.W., et al. Prenatal famine and genetic variation are independently and additively associated with DNA methylation at regulatory loci within IGF2/H19. PloS one 7, e37933 (2012).

20. Pidsley, R., Dempster, E.L. & Mill, J. Brain weight in males is correlated with DNA methylation at IGF2. Molecular psychiatry 15, 880–881 (2010).

## Methods References

1. Yu, P., McKinney, E.C., Kandasamy, M.M., Albert, A.L. & Meagher, R.B. Characterization of brain cell nuclei with decondensed chromatin. Dev Neurobiol 75, 738–756 (2015).

2. Matevossian, A. & Akbarian, S. Neuronal nuclei isolation from human postmortem brain tissue. J Vis Exp (2008).

3. Triche, T.J., Jr., Weisenberger, D.J., Van Den Berg, D., Laird, P.W. & Siegmund, K.D. Low-level processing of Illumina Infinium DNA Methylation BeadArrays. Nucleic acids research 41, e90 (2013).

4. McCartney, D.L., et al. Identification of polymorphic and off-target probe binding sites on the Illumina Infinium MethylationEPIC BeadChip. Genomics data 9, 22–24 (2016).

5. Ritchie, M.E., et al. limma powers differential expression analyses for RNA-sequencing and microarray studies. Nucleic acids research 43, e47 (2015).

6. Chang, C.C., et al. Second-generation PLINK: rising to the challenge of larger and richer datasets. GigaScience 4, 7 (2015).

7. Purcell, S., et al. PLINK: a tool set for whole-genome association and population-based linkage analyses. American journal of human genetics 81, 559–575 (2007).

8. Pedersen, B.S., Schwartz, D.A., Yang, I.V. & Kechris, K.J. Comb-p: software for combining, analyzing, grouping and correcting spatially correlated P-values. Bioinformatics (Oxford, England) 28, 2986–2988 (2012).

9. Roche. How to Evaluate SeqCap EZ Target Enrichment Data. (2017).

10. Bolger, A.M., Lohse, M. & Usadel, B. Trimmomatic: a flexible trimmer for Illumina sequence data. Bioinformatics (Oxford, England) 30, 2114–2120 (2014).

11. Xi, Y. & Li, W. BSMAP: whole genome bisulfite sequence MAPping program. BMC bioinformatics 10, 232 (2009).

12. Li, H., et al. The Sequence Alignment/Map format and SAMtools. Bioinformatics (Oxford, England) 25, 2078–2079 (2009).

13. Anderson, C.A., et al. Data quality control in genetic case-control association studies. Nature protocols 5, 1564–1573 (2010).

14. Altshuler, D.M., et al. Integrating common and rare genetic variation in diverse human populations. Nature 467, 52–58 (2010).

15. Das, S., et al. Next-generation genotype imputation service and methods. Nature genetics 48, 1284–1287 (2016).

16. Loh, P.R., Palamara, P.F. & Price, A.L. Fast and accurate long-range phasing in a UK Biobank cohort. Nature genetics 48, 811–816 (2016).

17. Genomes Project, C., et al. A global reference for human genetic variation. Nature 526, 68–74 (2015).

18. Biological insights from 108 schizophrenia-associated genetic loci. Nature 511, 421–427 (2014).

19. Won, H., et al. Chromosome conformation elucidates regulatory relationships in developing human brain. Nature 538, 523–527 (2016).

20. Dobin, A., et al. STAR: ultrafast universal RNA-seq aligner. Bioinformatics (Oxford, England) 29, 15–21 (2013).

21. Newman, A.M., et al. Robust enumeration of cell subsets from tissue expression profiles. Nature methods 12, 453–457 (2015).

22. Yu, Q. & He, Z. Comprehensive investigation of temporal and autism-associated cell type composition-dependent and independent gene expression changes in human brains. Scientific reports 7, 4121 (2017).

23. Darmanis, S., et al. A survey of human brain transcriptome diversity at the single cell level. Proceedings of the National Academy of Sciences of the United States of America 112, 7285–7290 (2015).

24. Croning, M.D., Marshall, M.C., McLaren, P., Armstrong, J.D. & Grant, S.G. G2Cdb: the Genes to Cognition database. Nucleic acids research 37, D846–851 (2009).

25. Harrow, J., et al. GENCODE: the reference human genome annotation for The ENCODE Project. Genome research 22, 1760–1774 (2012).

26. Subramanian, A., et al. Gene set enrichment analysis: a knowledge-based approach for interpreting genome-wide expression profiles. Proceedings of the National Academy of Sciences of the United States of America 102, 15545–15550 (2005).

27. Romero, P., et al. Computational prediction of human metabolic pathways from the complete human genome. Genome biology 6, R2 (2005).

28. Kandasamy, K., et al. NetPath: a public resource of curated signal transduction pathways. Genome biology 11, R3 (2010).

29. Croft, D., et al. The Reactome pathway knowledgebase. Nucleic acids research 42, D472–477 (2014).

30. Fabregat, A., et al. The Reactome pathway Knowledgebase. Nucleic acids research 44, D481–487 (2016).

31. Schaefer, C.F., et al. PID: the Pathway Interaction Database. Nucleic acids research 37, D674–679 (2009).

32. Mi, H., et al. The PANTHER database of protein families, subfamilies, functions and pathways. Nucleic acids research 33, D284–288 (2005).

33. Ashburner, M., et al. Gene ontology: tool for the unification of biology. The Gene Ontology Consortium. Nature genetics 25, 25–29 (2000).

34. Merico, D., Isserlin, R. & Bader, G.D. Visualizing gene-set enrichment results using the Cytoscape plug-in enrichment map. Methods in molecular biology (Clifton, N.J.) 781, 257–277 (2011).

35. Tang, S.H., Silva, F.J., Tsark, W.M. & Mann, J.R. A Cre/loxP-deleter transgenic line in mouse strain 129S1/SvImJ. Genesis 32, 199–202 (2002).

36. Ferron, S.R., et al. Differential genomic imprinting regulates paracrine and autocrine roles of IGF2 in mouse adult neurogenesis. Nature communications 6, 8265 (2015).

37. Mikaelsson, M.A., Constancia, M., Dent, C.L., Wilkinson, L.S. & Humby, T. Placental programming of anxiety in adulthood revealed by Igf2-null models. Nature communications 4, 2311 (2013).

38. Christian, R., Saavedra L, Gaynes BN, Sheitman B, Wines RCM, Jonas DE, Viswanathan M, Ellis AR, Woodel C, Carey TS. Future Research Needs for First- and Second-Generation Antipsychotics for Children and Young Adults (Agency for Healthcare Research and Quality, 2012).

39. Akbarian, S., et al. The PsychENCODE project. Nature neuroscience 18, 1707–1712 (2015).

40. Quinlan, A.R. & Hall, I.M. BEDTools: a flexible suite of utilities for comparing genomic features. Bioinformatics (Oxford, England) 26, 841–842 (2010).

41. Li, H. Tabix: fast retrieval of sequence features from generic TAB-delimited files. Bioinformatics (Oxford, England) 27, 718–719 (2011).

42. Robinson, M.D., McCarthy, D.J. & Smyth, G.K. edgeR: a Bioconductor package for differential expression analysis of digital gene expression data. Bioinformatics (Oxford, England) 26, 139–140 (2010).

43. Fortin, J.P., Triche, T.J., Jr. & Hansen, K.D. Preprocessing, normalization and integration of the Illumina HumanMethylationEPIC array with minfi. Bioinformatics (Oxford, England) (2016).

44. Shannon, P., et al. Cytoscape: a software environment for integrated models of biomolecular interaction networks. Genome research 13, 2498–2504 (2003).

45. Merico, D., Isserlin, R., Stueker, O., Emili, A. & Bader, G.D. Enrichment map: a network-based method for gene-set enrichment visualization and interpretation. PloS one 5, e13984 (2010).

46. Kucera, M., Isserlin, R., Arkhangorodsky, A. & Bader, G.D. AutoAnnotate: A Cytoscape app for summarizing networks with semantic annotations. F1000Research 5, 1717 (2016).

